# Massively parallel characterization of engineered transcript isoforms using direct RNA sequencing

**DOI:** 10.1101/2021.01.02.425091

**Authors:** Matthew J. Tarnowski, Thomas E. Gorochowski

## Abstract

Transcriptional terminators signal where transcribing RNA polymerases (RNAPs) should halt and disassociate from DNA. However, because termination is stochastic, two different forms of transcript could be produced: one ending at the terminator and the other reading through. An ability to control the abundance of these transcript isoforms would offer bioengineers a mechanism to regulate multi-gene constructs at the level of transcription. Here, we explore this possibility by repurposing terminators as ‘transcriptional valves’ which can tune the proportion of RNAP read-through. Using one-pot combinatorial DNA assembly we construct 1183 transcriptional valves for T7 RNAP and show how nanopore-based direct RNA sequencing (dRNA-seq) can be used to simultaneously characterize the entire pool at a nucleotide resolution *in vitro* and unravel genetic design principles to tune and insulate their function using nearby sequence context. This work provides new avenues for controlling transcription and demonstrates the value of long-read sequencing for exploring complex sequence-function landscapes.

## Introduction

The ability to precisely control when and where genes are expressed is crucial for engineering the behavior of living cells. To achieve this, bioengineers have predominantly focused on developing genetic parts to regulate the transcriptional and translational initiation rates of genes of interest ^1^. However, endogenous gene regulation is often multifaceted, employing diverse mechanisms that affect the stability and processing of DNA, RNA and proteins to create complex regulatory programs ^1,2^.

Transcript isoforms are commonly used by eukaryotes to diversify the products produced by a single gene and are generally created by the subsequent processing of a transcript by splicing machinery ^3^. Although such machinery is not generally present in prokaryotes, there is growing realization that these organisms also produce transcript isoforms by utilizing incomplete transcriptional termination ^4,5^. In this case, two transcript isoforms are possible: one ending at the terminator (if termination succeeds), and the other reading through (if termination fails) (**Figure 1A**).

**Figure 1:**
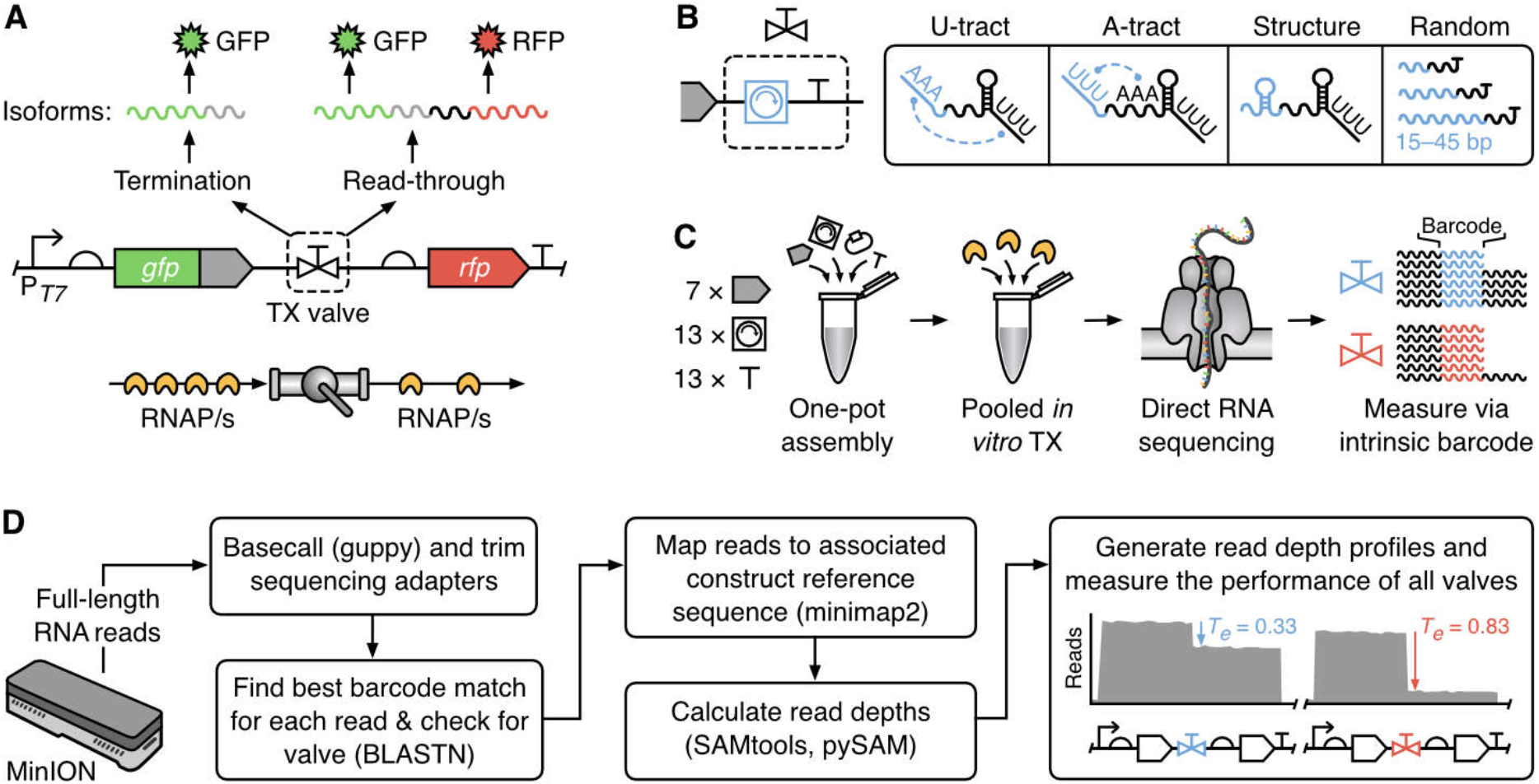
Massively parallel characterization of transcript isoforms using direct RNA sequencing. (**A**) Schematic of the genetic construct used to characterize transcriptional valves. A transcriptional valve controls the ratio of RNA polymerase (RNAP) termination to read-through and thus the proportions of transcript isoforms produced. (**B**) Our modular transcriptional valves comprise a ‘core terminator’ and ‘modifier’ sequence (blue) used to tune termination efficiency. Various modifiers were considered in this study to interact with the U- and A-tract of the core terminator, form small secondary structures in the RNA, and act as different length inert spacing elements. (**C**) The steps involved in the assembly of the modular transcriptional valve library and its pooled characterization using nanopore-based direct RNA sequencing. (**D**) Analysis pipeline used to generate valve specific read depth profiles and calculate termination efficiencies (*T_e_*) from pooled direct RNA sequencing data. Key computational tools shown in parentheses.

The design of synthetic terminators has so far focused on increasing their strength to achieve near perfect termination ^6^ as a way of insulating multiple genetic parts and devices from each other ^7^ and the host genome ^8^. Attempting to engineer terminators that function poorly in terms of efficiency or which can be tuned for manipulating transcript isoform ratios has not yet been explored. However, an ability to design transcript isoforms in this way could open up new avenues to control the stoichiometry of multi-gene expression purely at the level of transcription. This regulatory approach could be more efficient than other more commonly used methods, like operons, as not all RNA polymerases (RNAPs) would need to synthesize full length multi-gene transcripts and thus could be freed more quickly for other tasks. This approach would also be suitable for differential expression of purely RNA-based regulators (e.g. small RNA triggers for toehold switches ^9^ or gRNA arrays ^10^) where translation into protein does not occur.

A major bottleneck when developing new genetic parts is the time and effort needed to characterize libraries of parts to understand their design principles and build predictive models of their function. For transcriptional terminators a commonly used approach *in vivo* is a fluorescence assay in which two different fluorescent reporter proteins have the terminator to be tested placed between them. By comparing the ratio of fluorescence for each reporter with and without the terminator present, it is possible to quantify the fraction of transcriptional read through and the termination efficiency of the terminator ^6,11,12^. More recently, methods employing RNA sequencing (RNA-seq) have been used to provide a more detailed and direct measurement of termination at a nucleotide resolution by allowing for transcriptional profiles capturing RNAP flux along the DNA to be inferred. Drops in RNAP flux within these profiles can be used to measure both termination efficiencies as well as the precise location where these events occur ^13,14^. A challenge with all *in vivo* approaches is that they often need to make assumptions about the stability of the cellular environment and properties of the transcripts and proteins produced. However, these may not always hold: variations in mRNA stability ^15^, the occlusion of adjacent ribosome binding sites due to mRNA secondary structures, the impact of a terminator on translational coupling of neighboring genes ^6^, and transcription-translation coupling ^12^ all potentially playing a role for some designs and affecting the accuracy of the measurements made. Such differences may explain why *in vitro* and *in vivo* termination efficiencies have not been found to correlate well ^16^. Nonetheless, insights into *in vitro* termination have proven useful. Mairhofer *et al.* measured termination *in vitro* using chip-based capillary electrophoresis ^17^, identifying novel terminators which have subsequently been used both *in vitro* ^18^ and *in vivo* ^19^. A bigger issue facing all of these approaches though is their limited throughput and scalability with designs needing to be tested separately. This severely hampers the exploration of the vast genetic design space and limits our ability to understand how terminators are effectively engineered.

In this work, we aim to address these issues and develop a novel means for regulating multi-gene constructs. First, rather than treating terminators as the end point of a transcript, we consider them as ‘valves’ able to regulate the RNAP flux passing through a point in a DNA construct and thus the ratio of transcript isoforms that occur (**Figure 1A**). A transcriptional valve is characterized by its termination efficiency, calculated here as the probability that an RNAP terminates at the valve rather than continuing to transcriptionally read through. By designing transcriptional valves with varying efficiencies and placing them between genes of interest, ratios of RNAP flux, and thus the transcription rates of multiple genes, can be coupled to the activity of a single input promoter. This is similar to how an operon allows for robust control of protein expression stoichiometries due to varying translation rates of several genes encoded by a single transcript. To explore this approach, we design a library of 1183 modular transcriptional valves for T7 RNAP. The use of T7 RNAP ensures that the parts developed can be used in both cell-free systems and broadly across organisms in the future. We then show how nanopore-based direct RNA sequencing (dRNA-seq) can be used to characterize the *in vitro* function of the entire pooled library at a nucleotide resolution. Using this data, we are able to infer design principles and show how genetic context can be used to ‘tune’ termination efficiency or insulate a valve’s performance from local variable sequences. Our characterization methodology and experimental findings offer a novel means to control RNAP flux and transcript isoforms in genetic circuits and demonstrate how long-read sequencing can improve our understanding of large genetic design libraries.

## Results

### Designing transcriptional valves to control transcript isoforms

To demonstrate how transcriptional valves might be built, we attempted to construct proof-of-concept designs for T7 RNAP. T7 RNAP was selected due to its broad use in synthetic biology, which stems from the fact that it is a single-subunit RNAP with high processivity, making it ideal for both *in vitro* use ^20^ as well as an orthogonal transcription system *in vivo* ^21,22^. Whilst diverse terminators are available for native *Escherichia coli* RNAP ^6,11^, for T7 RNAP only a single terminator exists in the T7 phage genome ^23^ and only a few alternatives have been characterized ^17,24,25^. Termination of RNAP in model microorganisms (e.g. *E. coli* and *S. cerevisiae*) has also been extensively studied ^26^. However, T7 RNAP termination has many unknowns such as alternative intrinsic terminators beyond those in the T7 phage genome and the bidirectionality of termination.

Specific features of a terminator, such as its hairpin structure and U-tract, can strongly influence termination efficiency ^6,27^. Therefore, as a basis for a library of transcriptional valves, we chose 13 different intrinsic terminators to act as ‘core terminating’ elements (T). We began by selecting the single terminator from the T7 phage genome (T27) ^23^, which has previously been characterized *in vitro* ^17^. To test its possible bidirectionality, a feature that terminators for other RNAPs have been shown to exhibit ^6^, it was included in a reverse orientation in our designs ^17^. Beyond native phage terminators, *E. coli* terminators present another source of these parts and have been shown to terminate T7 RNAP *in vitro ^17,28^*. Therefore, 11 intrinsic rho-independent terminators were selected from the *E. coli* genome spanning a wide range of termination efficiencies *in vivo ^6^*. Finally, a negative control terminator (T33) was designed. This consisted of a random non-coding sequence generated using R2oDNA designer ^29^ and was verified to not contain a hairpin in the mRNA secondary structure using the viennaRNA tool ^30^.

The genetic sequence immediately upstream of a terminator-hairpin also influences termination ^11,12^ and we reasoned that this region could be used to fine-tune the termination efficiency of a valve. Therefore, we included a ‘modifier’ part (M) in our transcriptional valve design and developed 13 different modifier sequences (**Figure 1B**). Our first set of modifiers contained motifs designed to interact with canonical regions of a terminator hairpin sequence. Specifically, modifiers M10 and M11 were designed to interact with possible U- and A-tracts within a terminator by containing complementary homopolymers of adenine or uracil, respectively ^6^ (**Figure 1B**). A further modifier M13 was designed to encode a small RNA secondary structure with the goal of affecting RNA structure formation near the terminator part. Beyond tuning termination efficiency with RNA interactions and structures, it has been shown that inert random sequences can play an insulating role, improving the robustness of a genetic part’s performance ^31^. It is also known that upstream genetic context of intrinsic terminators influences termination in a distant-dependent manner ^11,12^. We therefore decided to include a selection of modifiers of different lengths (M13–M16: 15 bp, M17–M19: 30 bp and M20–M22: 45 bp) where each was a random non-coding sequence generated by R2oDNA designer ^29^.

To assess the robustness of each valves’ termination efficiency to local upstream genetic context, our library also included ‘spacer’ elements (S) (**Figure 1B**). These did not form part of the transcriptional valve, but instead allowed us to see how a particular valve might behave when used in combination with other components (e.g. protein coding regions). Using the NullSeq tool ^32^, we generated 7 random and genetically diverse 33 bp long spacers with a nucleotide composition similar to coding regions of *E. coli* that could be placed at the 5’ end of a valve. Each spacer had a stop codon ‘TAA’ at its 3’-end though this was not utilized in our *in vitro* assay. Taken together, our spacers, modifiers and terminators could be combinatorically assembled to create a total of 1183 unique designs able to regulate RNAP flux and provide valuable information regarding the design principles of transcriptional valves.

### Combinatorial assembly of a transcriptional valve library

A one-pot pooled combinatorial DNA assembly method was used to physically construct the final set of transcriptional valve designs (**Figure 1C**, **Methods**) ^33^. To act as a backbone for controlling expression of our constructs and to enable their maintenance and propagation in *E. coli* cells, we modified a plasmid which had previously been used to characterize terminators *in vivo* (pGR) ^6^. The pGR plasmid consists of an *araC* expression cassette and arabinose inducible promoter (P_*BAD*_) that drives expression of a *gfp* reporter gene. Directly downstream of the *gfp* expression cassette is an *rfp* reporter gene that is expressed only if transcriptional readthrough from *gfp* occurs. Initially, no transcriptional terminator is included between these genes and typically the small spacer region that is present is replaced with the terminator to be characterized. The plasmid also includes an ampicillin selection marker and a ColE1 origin of replication leading to a copy number of ~15–20 per cell. To make this plasmid compatible with T7 RNAP, we replaced the *araC* expression cassette and P_*BAD*_ promoter with a strong constitutive T7 RNAP promoter (P_*T7*_) and removed the stop codon at the end of the *gfp* gene to allow the spacer region to be appended directly to the end of the coding sequence (**Figure 1A**, **Methods**).

DNA assembly was performed by carrying out a dual digestion of the modified pGR plasmid to cut the plasmid at two points between the *gfp* and *rfp* genes. This resulted in linearized plasmid DNA with 4 nucleotide (nt) 3’ overhangs that enabled insertion of the parts necessary to assemble transcriptional valves. To reduce costs, the spacer, modifier and terminator elements were synthesized as single-stranded forward and reverse DNA oligonucleotides that when annealed together created double stranded DNA fragments with 4 nt 5’ overhangs on each end. Overhangs were designed to hybridize specifically to the overhangs of DNA fragments encoding adjacent parts or the digested plasmid backbone (**Methods**). Overhangs were selected based on experimental evidence showing a lack of cross-reactivity ^6,34^. By mixing equimolar amounts of each type of DNA part in a single DNA ligation reaction, the entire combinatorial library was assembled. *E. coli* cells were then transformed with this pooled library and approximately 500,000 colonies (>400-fold library coverage) selected from plates via scraping before their pooled DNA was extracted. Such a high fold-coverage guaranteed representation of each design in the sample (**Supplementary Note 1**) ^35^.

Nanopore-based long-read DNA sequencing (DNA-seq) was then used to verify the successful combinatorial assembly of all 1183 genetic constructs. This showed that all designs were present with an even distribution of parts but an uneven distribution of designs (**Supplementary Figure 1**). Part frequencies were similar to ratios expected from equiprobable assembly, except for short parts 15 bp long, which may be due to reduced efficiency during the combinatorial assembly. These part frequencies were used to predict design frequencies, of which 91 had >20% absolute deviation between predicted and measured frequency. Certain parts were overrepresented in these 91 designs (M14: 3-fold and T18: 2.3-fold) indicating that the abundance of constituent parts does not solely dictate design abundance within the library.

Finally, we investigated DNA assembly fidelity by generating accurate consensus sequences from the long-read DNA-seq data (**Methods**). Comparing the reference and consensus sequence for each design we found a mean of 0.6 single nucleotide polymorphisms (SNPs) per design, with 40% of designs having no SNPs (**Supplementary Figure 1**). Using the same analysis, we found that no designs had a SNP for an equivalent (100 bp) length region of the GFP encoding sequence.

### Pooled characterization using direct RNA sequencing

It would be challenging to characterize this large library using typical fluorescence-based methods because each construct would need to be isolated from the pool and separately assayed ^6^. The use of RNA sequencing (RNA-seq) has emerged as an alternative method able to provide detailed nucleotide resolution data related to RNAP flux that can be used to characterize genetic parts ^13,14,36^. However, existing RNA-seq approaches also suffer from the need to isolate and individually barcode RNA produced by each design.

To overcome this issue, we used nanopore based direct RNA sequencing (dRNA-seq) to provide full-length reads of transcript sequences ^37^. Crucially, each transcript isoform encodes its associated transcriptional value sequence either at the 5’ end, if termination was successful, or within the body of the transcript, if transcriptional read-through had occurred – the design’s sequence therefore acts as an ‘intrinsic barcode’. This allowed for individual reads to be attributed to a particular design without the need to separate them before preparing the sequencing library. Such an approach is not possible when using more common short-read-based RNA-seq because the transcriptional valve sequence for reads associated with read-through events will not always be located within 250 bp (the typical maximum read length for Illumina sequencing) of the transcript end, and so would not be captured. Instead, by using long-read dRNA-seq the whole library can be assayed as a single pooled sample and the data demultiplexed to produce separate read depth profiles for each design. By comparing the ratio of transcript isoforms for each design (i.e. read depth directly before and after the transcriptional valve) a termination efficiency can be calculated (**Figure 1C**).

While this approach removes the need to separate and attach unique ‘barcode’ sequences to each design/sample when characterizing the library, it does rely on each transcriptional valve having a sufficiently different sequence for each read to be accurately mapped to a single design. This limits the detail of the genetic design space that can be explored. Read accuracy for nanopore-based dRNA-seq is, at present, lower than for Illumina-based short-read RNA-seq (median read accuracy of 80-90% versus >99.9%, respectively ^38^), although this gap is closing with improvements to basecallers and sequencing chemistries. Therefore, analysis pipelines need to be carefully tuned and validated to ensure accurate demultiplexing of reads when dRNA-seq is used in this way.

### Optimizing the computational analysis pipeline

To optimize the analysis pipeline and ensure that our library could be accurately characterized, we developed a simple computational model to simulate the error-ridden reads that would be obtained after dRNA-seq. In our simulations, errors took the form of random nucleotide substitutions. At present, dRNA-seq reads have typical accuracies of 85–95% and so we selected a 15% substitution frequency in our model. Whilst other types of error such as insertions, deletions and elevated error rates at homopolymers were not included in our model, we found that our simulations were sufficient to identify key parameters and part dissimilarity thresholds for effective demultiplexing of long sequencing reads.

To demultiplex the dRNA-seq reads, the BLASTN tool was used to find all possible alignments between a read and the library of designs, with the best match being chosen ^39,40^. Optimizing the BLASTN parameters is crucial for accurate characterization and so computational experiments were performed where a smaller library (540 designs, **Supplementary Figure 2**) was used to systematically explore the role of each BLASTN parameter (**Methods**). Each design was given a random termination efficiency and had a set of full-length terminated and non-terminated reads generated based on our model. These reads were then pooled for all the designs and attempts made to demultiplex and infer the original termination efficiencies for each design. We found that the key BLASTN parameters were: ‘word size’ which was reduced from its default of 11 to the minimum of 4, and the ‘reward for a nucleotide match’ which was raised from its default of 1 to 2.

The final computational demultiplexing and analysis pipeline (**Figure 1D**) involved aligning the sequences of all designs against all reads using BLASTN with optimized parameters and then associating each read with the design that had the best alignment score (**Methods**). An e-value threshold of 1 was selected such that only alignments where one or fewer hits of a similar score would be expected to be seen by chance from a sequence database of that size. Reads for each design were then mapped to the appropriate reference sequence using minimap2 ^39^ and design-specific read depth profiles generated using SAMtools and pySAM ^41^filtering out any reads where no terminator was present after alignment and mapping. Finally, termination efficiencies were calculated for each read depth profile as *T_e_* = [*R*(*x_s_) – R*(*x_e_*)] / *R*(*x_s_)*, where *R*(*x*) is the read depth at position *x* in the genetic construct, and *x_s_* and *x_e_* are the start and end position of the transcriptional valve, respectively.

### In vitro characterization

*In vitro* transcription using T7 RNA polymerase of the entire pool of DNA constructs followed by dRNA-seq enabled us to rapidly assay the performance of each design concurrently (**Figure 1C**). To ensure the accuracy of our measurements, a detailed analysis of the generated read depth profiles was performed. This revealed several key features (**Figure 2**). First, we noticed that dRNA-seq reads often had 6 nt of their 5’ sequence truncated (**Figure 2A**), which could make it difficult to determine precise transcription start sites. As dRNA-seq progresses from the 3’ to 5’ end of an RNA molecule, this short region likely corresponds to the point where the motor protein that ratchets the RNA molecule through the pore reaches the 3’ end of the molecule and releases it, causing an increased error or removal of the short sequence contained within the pore.

**Figure 2.**
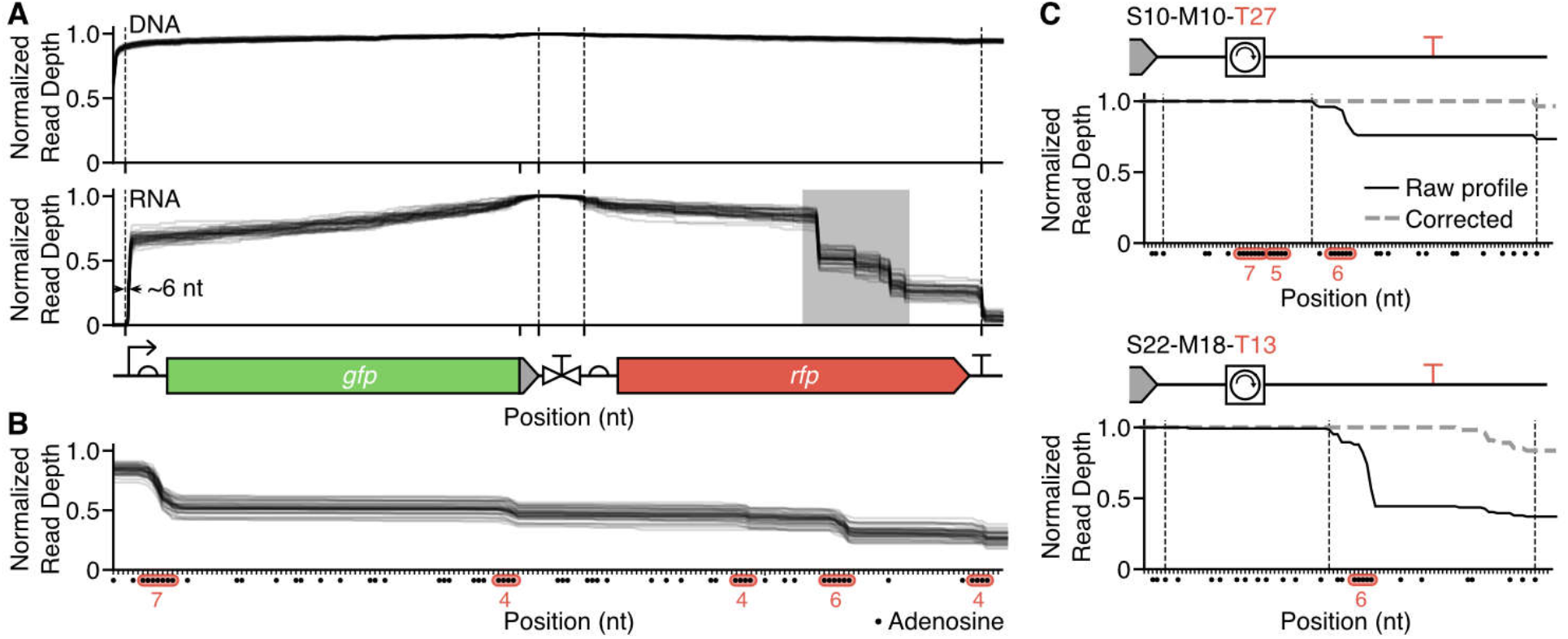
Features of direct RNA sequencing read depth profiles. (**A**) Normalized sequencing read depth profiles from nanopore-based DNA-seq and dRNA-seq for 42 designs containing the same core terminator T33 (non-terminating control) and modifiers of length 30 nucleotides. Vertical dotted lines denote transcript and valve boundaries. Plasmid map illustrated beneath, to scale. Grey shaded region is expanded in panel B. (**B**) Expanded region from panel A showing dRNA-seq read depth profiles with dots corresponding to adenosine nucleotides. Adenosine homopolymers ≥4 nt in length are highlighted in red and their lengths are shown below. (**C**) Corrected (dashed grey lines) and raw (solid black lines) dRNA-seq read depth profiles for two different designs where the core terminator contains an adenosine homopolymer within the terminator sequence. Vertical dotted lines indicate spacer-modifier and modifier-terminator boundaries.

Second, we found that all dRNA-seq read depth profiles showed drops in read depth when moving away from the intrinsic barcode sequence (**Figure 2A**). Such a feature is found in all nanopore dRNA-seq studies to date covering RNA samples from many different organisms ^37,38^ and is thought to arise due to fragmentation of full-length RNA molecules (e.g. by shearing caused during pipetting) and/or premature abortion of sequencing single molecules. In contrast, only small drops were observed for nanopore DNA sequencing of the constructs (**Figure 2A**), possibly due to the greater stability of the molecule ^42^.

To validate this hypothesis and explore its possible impact on *T_e_* measurements, we developed a mathematical model (**Supplementary Note 2**) and used data from an RNA Control Strand (RNA CS) that is externally ‘spiked-in’ to each dRNA-seq experiment for quality control assessments. Because the RNA CS is a single fixed length sequence, we could use it to test how different amounts of fragmentation or sequencing termination affect the shape of a read depth profile recovered. We found that the experimental data could be well described by a simple model with three probabilistic processes: fragmentation before ligation of sequencing adapters, successful adaptor ligation, and sequencing read truncation (**Supplementary Note 2**). Sequencing read truncation could be caused by RNA fragmentation (after adapter ligation) and/or early abortion of the sequencing process. We found that the impact of these effects on *T_e_* was small (**Supplementary Figures 3 and 4**), however, fragmentation did cause a significant reduction in the number of reads mapping to an intrinsic barcode (~20% of the total reads mapped to a barcode). Therefore, improvements in experimental protocols to reduce fragmentation or the incorporation of methods to enrich barcode containing reads (e.g. using ‘read until’ technologies ^43^ or sequence-specific dRNA-seq) could both improve the accuracy of *T_e_* calculations and increase the size of design libraries that can be assessed using a single sequencing run.

A third observation was that significant drops in read depth were seen outside of the core terminators and predominantly at short poly-A sequences >4 nt in length (**Figure 2B**). When preparing RNA for dRNA-seq a poly-A tail is required for ligation of sequencing adapters to the 3’-end of the RNA molecules. As *in vitro* transcription of our constructs will not produce transcripts of this form, we used *E. coli* poly(A) polymerase to polyadenylate all the RNAs produced (**Methods**). Analysis of the dRNA-seq data showed <10 nt poly-A tails were present, which were shorter than other dRNA-seq runs we had previously performed (**Supplementary Figure 5**). We hypothesized that inefficient polyadenylation allows for fragmented RNAs with a short poly-A end to become enriched during sequencing and thus causes notable drops at these points within a construct that do not correspond to termination events. This was supported by another dRNA-seq run where efficient polyadenylation resulted in the removal of these drops. This feature only affects the measurement of *T_e_* for designs containing parts with poly-A regions in their template strand (i.e. I10, T13 and T27) and is easily mitigated by retaining only mapped reads which do not terminate at a poly-A motif outside the terminator hairpin (**Figure 2C**).

After correcting for these effects, we found that transcript abundances correlated with DNA construct frequencies (*R*^2^ = 0.48; **Supplementary Figure 1**) and a strong correlation (*R*^2^ = 0.84) was observed for calculated *T_e_* values between biological replicates (**Supplementary Figure 6**). Overall, *T_e_* varied from 0 to 0.73 across the library (**Figure 3A,B**) with some modifiers significantly affecting median *T_e_* in a systematic way (e.g. the positive influence of the U-tract and A-tract modifiers; **Figure 3C**) and core terminators displayed varying levels of sensitivity to different modifier and spacer parts (**Figures 3D**). Grouping designs by their core terminator, we found that the non-terminator part (T33) showed little to no termination (*T_e_* < 0.05) across the majority of designs. Similarly, for the reverse oriented T7 phage terminator (T27) all designs had *T_e_* < 0.05, indicating that it is not able to efficiently terminate T7 RNAP bidirectionally. The eleven *E. coli* terminators showed a range of termination efficiencies for T7 RNAP with median *T_e_* varying from 0.01 to 0.51. Four of the *E. coli* terminators (T15, T21, T17, T18) had exceptionally low efficiency, with 99% of their designs having *T_e_* < 0.05. The remaining seven *E. coli* terminators displayed a range of termination efficiencies heavily influenced by upstream sequence context.

**Figure 3.**
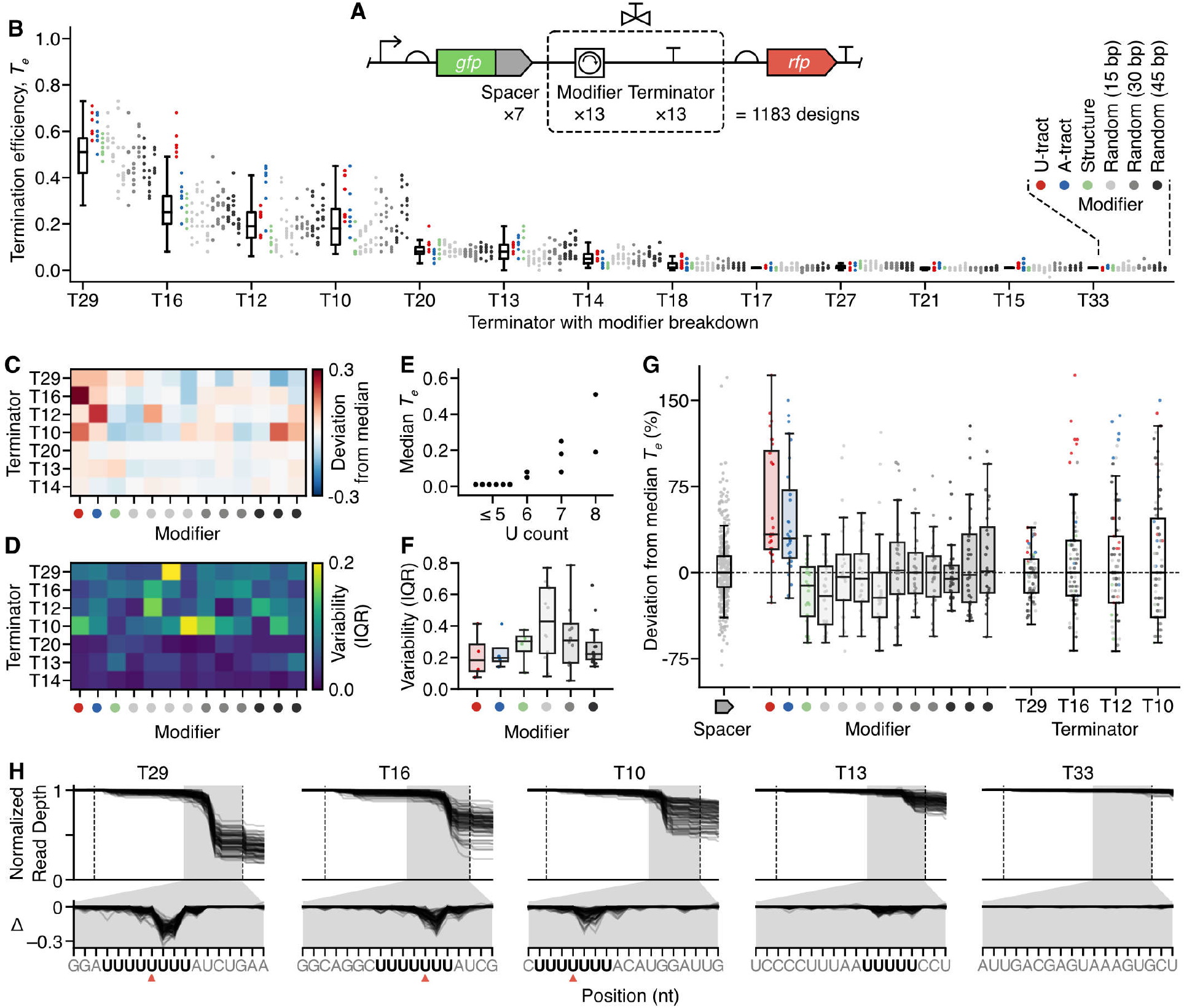
Characterization of a T7 RNAP transcriptional valve library. **(A)** Structure of the pooled transcriptional valve library. Grey ‘spacer’ elements are 33 bp random sequences used to assess the sensitivity of a transcriptional valve to nearby sequence context. (**B**) Termination efficiency (*T_e_*) for every design in the library. Each point denotes the *T_e_* valve for a unique genetic construct color coded by the modifier present. Points are grouped by core terminator with a box plot summarizing the data for all associated constructs. (**C**) Deviation of *T_e_* from the median valve of all constructs containing the associated core terminator. Each square shows the median *T_e_* value for all constructs containing a specific core terminator and modifier. Only data for core terminators with median *T_e_* > 0.05 is shown. (**D**) Interquartile range (IQR) of *T_e_* values for all constructs containing a specific core terminator and modifier. Only data for core terminators with median *T_e_* > 0.05 is shown. (**E**) Median *T_e_* for designs containing each core terminator compared to the number of uracil (U) residues in the 8 nt from where the U-tract begins. (**F**) IQR of *T_e_* values for transcriptional valves grouped by modifier type. Only data for designs containing the strongest terminators (T29, T16, T12, T10) was used. (**G**) Percentage deviation in *T_e_* for each construct to the median of all designs containing the same core terminator. Each point corresponds to a single construct grouped by spacer, modifier type or core terminator. Only data for designs containing the strongest terminators (T29, T16, T12, T10) was used. (**H**) Normalized dRNA-seq read depth profiles for terminators and non-terminator control (T33). Each line corresponds to a design. Dotted lines denote the start and end of the core terminator. Grey shaded region is expanded in the lower panel showing the change in normalized read depth at each nucleotide position with respect to the previous nucleotide (Δ).

We assessed several predictors of termination efficiency and found the length of the core terminator U-tract to be the key determinant for T7 RNAP. Median *T_e_* values for designs with a common core terminator correlated with the number of U residues in the U-enriched 8 nt sequence included within or immediately downstream of the terminator hairpin (**Figure 3E**). The six core terminators with 5 or fewer U-residues had low termination efficiency. This matches a previous finding for *E. coli* RNAP termination ^6^. Other parameters explored included GC content and minimum free folding energy, none of which correlated with measured *T_e_* (**Supplementary Figure 7**).

### Insulating transcriptional valves from local genetic context

It has been shown for other types of genetic part that more reliable performance can also be obtained by inserting random non-coding sequencers around a part to insulate its function from potential interactions with other nearby part sequences ^7,31^. Our library specifically included random non-coding modifiers of varying length to assess whether similar insulating effects would be seen for transcriptional valves (**Figure 1B**). In general, we found that an increase in the length of these modifiers lead to a marked reduction in *T_e_* variability when an identical valve design was used across numerous genetic contexts (i.e. upstream spacer sequences; **Figure 3F**). This matches findings for bacterial promoters where longer upstream insulating sequences resulted in more predictable gene expression ^31^. Valves containing U-tract, A-tract and secondary structure modifiers also acted as effective insulators with low variability in *T_e_* across genetic contexts compared to equal length random modifiers. Notably, these effects were also terminator specific, with some core terminators showing more predictable behavior across modifiers (e.g. T29) than others (e.g. T10) (**Figure 3G**). This suggests that some terminators might be better suited to tuning, whilst others are better placed to maintain a consistent termination efficiency.

### Tuning the strength of transcriptional valves

It is known that local sequence context can affect the function of genetic parts. We therefore designed modifiers with the aim of being able to tune the strength of a valve. We found that changes in upstream genetic context (i.e. both modifier and spacer sequences) could significantly influence termination strength, allowing *T_e_* to be varied over a range of 0.6 for designs containing the stronger core terminators such as T16 (**Figure 3G**). Notably, modifiers within the valves were often found to have a similar tuning effect across many different core terminators. For example, the U- and A-tract modifiers enhanced the termination efficiency of most designs (**Figure 3G**). However, large variability in effect sizes was seen across the different valves suggesting that sequence specific features also play a key role in modulating the precise termination efficiencies achieved. We found that spacers did not have such a systematic impact. For designs grouped by spacer, the median percentage deviation from the median of the valve they contained was found to be <5% (**Figure 3G**), suggesting that tuning of termination efficiency is best achieved by varying sequence context close to the core terminator part.

### Nucleotide resolution insights

A useful feature of genetic part characterization by RNA-seq is the ability to extract nucleotide resolution insights from the read depth profiles. To enable comparisons between our designs where total numbers of reads for each design varied, we generated profiles normalized by the read depth at the start of the transcriptional valve such that drops corresponded to a fractional change (**Figure 3H**). We also calculated Δ-values corresponding to the change in normalized read depth at each nucleotide position with respect to the previous nucleotide, enabling us to pinpoint and compare changes more easily.

For all valves, drops in read depth occurred as expected in the U-tract with little variation in position seen between designs. Moreover, the major drops in the profiles occurred at a similar position (after 4 uridines) in the U-tract for the stronger core-terminators (**Figure 3H**), which likely captures a point where a combination of T7 RNAP pausing and the stability of the transcription elongation complex is sufficiently weakened to cause termination. The ability to observe these nucleotide resolution changes offers another means to characterize the functionality of genetic parts (e.g. the precise locations where termination occurs) and demonstrates a further benefit of the pooled dRNA-seq over other methods.

## Discussion

In this work, we have shown how transcriptional terminators can be repurposed as ‘valves’ to regulate the flow of RNAP along DNA and control the ratio of transcript isoforms produced. By developing a new nanopore-based dRNA-seq characterization approach (**Figure 1**), we were able to simultaneously measure the termination efficiency of an entire mixed pool of 1183 unique transcriptional valves as well as provide nucleotide resolution insight into precisely where termination occurred for each (**Figure 3**). Such detail is lost with more typical fluorescence-based assays ^6,11^, but is essential for developing the low-level biophysical or machine learning based models of genetic parts that are essential for predictive biodesign workflows ^44–47^. While rich, high-content characterization data can normally only be produced for a small set of samples ^8,13,36,48^, the approach presented here removes this limitation, allowing us to more systematically explore the genetic design space of a large pooled library and glean several design principles. Specifically, local sequence context (i.e. modifier sub-sequences) can be used to tune termination efficiency, while the inclusion of sufficiently long insulating sequences at the 5’ end of a core terminator strongly reduce changes in *T_e_* when the same valve is used in conjunction with different upstream coding sequences. Furthermore, the successful use of *E. coli* terminators to control the viral T7 RNAP suggests that a similar characterization approach could be used to rapidly develop libraries of transcriptional valves for RNAPs from other organisms.

Although dRNA-seq opens up new avenues for high-throughput characterization of genetic part libraries, some challenges remain. The most prominent of these is ensuring the read depth profiles accurately represent the transcript variants present. Here, we show how unwanted features in the read depth profiles caused by the sequencing library preparation can be effectively corrected (**Figure 2**). However, improvements in the polyadenylation of transcripts or alternative methods for efficiently attaching sequencing adapters to transcripts to reduce these unwanted features would further refine the approach.

This work views transcriptional terminators in a new light. Not merely as a hard end point when producing a transcript, but as a means to tune and orchestrate one of the many flows that underpin the synthesis of proteins from DNA (e.g. transcription and translation). Nature is known to regulate gene expression at multiple levels and through numerous processes to create complex regulatory programs ^49^. Transcriptional valves offer bioengineers a new perspective on how multi-gene regulation can be implemented at a purely transcriptional level and a means to implement more diverse information flows in genetic circuitry.

## Methods

### Strains and media

All cloning was performed using *Escherichia coli* strain DH10-β (F^−^ *endA*1 *glnV*44 thi-1 *recA*1 *relA*1 *gyrA*96 *deoR nupG purB*20 φ80d*lacZ*ΔM15 Δ(*lacZYA*–*argF*)U169, *hsdR*17(r_K_^−^m_K_^+^), λ^−^-) (C3019I, New England Biolabs). Cells were grown in LB Miller broth (L3522, Sigma-Aldrich). Antibiotic selection was performed using 100 μg/mL of ampicillin (A9393, Sigma-Aldrich).

### Pooled combinatorial assembly of the transcriptional valve library

The pGR plasmid backbone ^6^ was modified for use in combinatorial assembly by first mutating the *gfp* stop codons from ‘TAATAA’ to ‘TTAGCA’ using Q5 mutagenesis (E0554S, New England Biolabs) and second replacing the *araC* gene and P_*BAD*_ promoter with a consensus T7 promoter sequence (pT7, **Supplementary Table 1**). Promoter substitution used the restriction enzymes AatII (10 units; R0117S, New England Biolabs) and NheI-HF (10 units; R3131S, New England Biolabs) in 1X CutSmart buffer (B7204S, New England Biolabs) and nuclease free water (final volume 50 μL) at 37 °C for 30 minutes; 20 minutes at 80 °C followed by adding annealed inserts to vector DNA (0.020 pmol) at a 3:1 molar ratio with T4 DNA ligase (400 units; M0202S, New England Biolabs), T4 DNA ligase buffer and nuclease free water to a final volume of 20 μL for 30 minutes at room temperature and then 65 °C for 10 minutes.

Oligonucleotides (25 nmol, dry lyophilized solid, standard desalting; Integrated DNA Technologies) were diluted to 100 μM in TE buffer (10 mM tris(hydroxymethyl)aminomethane, 0.1 mM ethylenediaminetetraacetic acid, pH 8.0). Duplex DNA was assembled for each core terminator, modifier, or spacer variant by annealing the complementary forward and reverse oligonucleotides (2 μL, 100 μM) in 46 μL annealing buffer (10 mM Tris, pH 7.5-8.0, 50 mM NaCL, 1 mM EDTA), heating to 95 °C for 5 minutes and slowly cooling to room temperature. Duplex DNA of all variants (1 μL (1 pmol) of each) was combined, and the pool was diluted to a final concentration of 1 pmol/μL with nuclease-free water. The pooled duplex DNA (20 μL) was phosphorylated using T4 polynucleotide kinase (10 units; M0201S, New England Biolabs) in 10X T4 DNA ligase buffer (2 μL) at 37 °C for 30 minutes; 65 °C for 20 minutes. Meanwhile, plasmid backbone DNA (1 μg) was digested using EcoRI-HF (20 units; R3101S, New England Biolabs) and SpeI-HF (20 units; R3133S, New England Biolabs) in 10X CutSmart buffer (5 μL) and nuclease-free water (35 μL) at 37 °C for 4 hours and then 80 °C for 20 minutes, before gel extraction (0.8 % agarose gel, gel green dye, 80V, 100 minutes; T1020S, New England Biolabs Monarch). Plasmid backbone (50 fmol) was used for pooled ligation based combinatorial assembly by combining with the aforementioned phosphorylated duplex DNA pool (containing 200 fmol of each part variant and therefore sufficient to make 200 fmol of each design), nuclease-free water (40 μL), 10X T4 DNA ligase buffer (5 uL) and T4 DNA ligase (320 units; M0202S, New England Biolabs) and incubating at room temperature for 3 hours and then 65 °C for 10 minutes.

The pooled transcriptional valve library (3 μL) was then added to each of 12 aliquots of *E. coli* strain DH10-β cells (45 μL each; C3019I, New England Biolabs) thawed on ice for 10 minutes and mixed gently by tapping. The mixture was left on ice for 30 minutes and heat-shocked at 42 °C for 30 seconds before leaving for 5 minutes on ice. NEB 10-beta/Stable Outgrowth Medium (450 μL; B9035S, New England Biolabs) was added to each aliquot and the mixture was shaken vigorously (1250 rpm) at 37 °C for 60 minutes. Each aliquot was then added to one 1.5 L rectangular glass dish (Pyrex) containing LB agar with ampicillin and cells were grown overnight. Following this, sufficient colonies were scraped from agar plates for >400-fold library coverage and DNA was extracted (T1010L, New England Biolabs). DNA concentrations were measured using a NanoPhotometer N60 (Implen).

### Verification of assembled transcriptional valve library

DNA from the pooled transcriptional valve library (400 ng) was prepared for DNA sequencing using the rapid barcoding kit following the standard protocol (SQK-RBK004, Oxford Nanopore Technologies). DNA samples were sequenced for 48 hours on FLO-MIN106 flow cells. Generated FAST5 files were basecalled using guppy version 3.1.5. BLASTN version 2.2.31 (with the same parameters as for dRNA-seq demultiplexing) was used to align sequencing reads to reference sequences. DNA sequencing reads were demultiplexed by selecting the best alignment to a design based upon maximum bitscore for each sequencing read. Sequencing reads with no alignment to a design, or alignments to multiple designs with the same maximum bitscore were excluded from further analysis. Part and design frequencies were calculated relative to the total number of annotated sequencing reads.

We used demultiplexed sequencing reads to generate a consensus sequence for each design and assess DNA assembly fidelity. Demultiplexed sequencing reads for each design were aligned to the plasmid encoding the design using minimap2 version 2.17 ^39^. Racon version 1.4 ^50^ with parameters: -m 8 -x -6 -g −8 -w 500, was used to polish the plasmid sequence and refine the consensus sequence produced. The polished and reference sequences were then aligned using Multiple Alignment using Fast Fourier Transform (MAFFT) with parameters: --localpair --maxiterate 1000. Finally, a custom Python script was used to count the average number of single nucleotide polymorphisms per design.

### Pooled in vitro transcription and direct RNA sequencing

DNA from the pooled transcriptional valve library (1 ug) was linearized using AatII (10 units) in 10X CutSmart buffer (5 μL) and nuclease-free water (40 μL) at 37°C for 30 minutes and then 80°C for 20 minutes. Duplicate reactions were purified (T1030L, New England Biolabs) and eluted in nuclease-free water (12 μL). *In vitro* transcription was performed using HiScribe™ T7 High Yield RNA Synthesis Kit (E2040S, New England Biolabs). The following reagents were combined: T7 RNA Polymerase mix (2 μL), adenosine triphosphate (2 μL), guanosine triphosphate (2 μL), cytidine triphosphate (2 μL), uridine triphosphate (2 μL), kit reaction buffer (2 μL) and the linearized DNA pool (250 ng) at 37 °C for 35 minutes. Synthesized RNA was diluted 20-fold in nuclease-free water and purified (R1013, Zymo Research). The purified RNA (20 μg) was then polyadenylated using *E. coli* Poly(A) Polymerase (10 units; M0276S, New England Biolabs) with nuclease-free water (29 μL), 10X reaction buffer (4 μL), RNase inhibitor murine (0.5 μL; M0314S, New England Biolabs) and 4 μL adenosine triphosphate, ATP (10 mM) at 37 °C for 30 minutes. The reaction was stopped by proceeding to RNA purification with elution in 15 μL (R1013, Zymo Research). Polyadenylated RNA (1 μg) was prepared using the direct RNA sequencing kit following the standard protocol (SQK-RNA002, Oxford Nanopore Technologies) with inclusion of the reverse-transcription step. RNA samples were sequenced for 48 hours on FLO-MIN106 flow cells.

### Computational demultiplexing and analysis pipeline

First, dRNA-seq data in FAST5 format was basecalled using guppy version 3.1.5 in high-accuracy mode. Next, the design sequences (not including plasmid backbone) were aligned to all basecalled sequencing reads using BLASTN version 2.2.31 ^51^. BLASTN parameters were selected based upon simulated nanopore RNA-seq data: -outfmt 6 -gapopen 5 -gapextend 2 - reward 2 -penalty −3 -evalue 1 -word_size 4 -max_target_seqs 1000000 -max_hsps 1. Custom Python and Bash scripts were then used to match each sequencing read to a design based upon the best alignment (the alignment with maximum bitscore). Sequencing reads with no alignment to a design, or alignments to multiple designs with the same maximum bitscore were excluded from further analysis. Part and design frequencies were calculated relative to the total number of annotated sequencing reads.

Parsed reads were then mapped to a plasmid sequence encoding the appropriate design using minimap2 version 2.17 ^39^ to generate a sequence alignment (SAM) file. Using pySAM, the SAM file was refined by removing sequencing reads which met any of the following criteria: (1) sequencing reads terminating at >4 consecutive adenosine residues and (2) sequencing reads that did not contain a full design sequence and terminated between the start of the spacer and end of the modifier as defined by a general feature format (GFF) file for each plasmid sequence bearing a design. Sequencing reads terminating within 3 nucleotides of criteria (1) or (2) were also omitted. Termination efficiencies were calculated based upon the read depth either side of the valve (see main text for details) which was further corrected by subtracting the predicted *T_e_* deviation (**Supplementary Note 2**; **Supplementary Figure 4**). Reads from two identical sequencing runs were pooled to calculate final *T_e_* values.

### Computational tools and genetic design visualization

Computational analyses were executed using Python version 3.5. All genetic diagrams are shown using Synthetic Biology Open Language Visual (SBOL Visual) notation ^52^. SBOL Visual diagrams were generated using the DNAplotlib Python package version 1.0 ^53^ which were then annotated and composed with OmniGraffle version 7.9.2.

## Acknowledgements

M.J.T. was supported by the EPSRC/BBSRC Centre for Doctoral Training in Synthetic Biology grant EP/L016494/1. T.E.G. was supported by a Royal Society University Research Fellowship grant UF160357 and BrisSynBio, a BBSRC/EPSRC Synthetic Biology Research Centre grant BB/L01386X/1.

## Author Contributions

T.E.G. conceived of the study and supervised the work. M.J.T. designed the genetic constructs, performed the experiments and developed data analysis pipelines. M.J.T. and T.E.G. analyzed the data and wrote the manuscript.

## References

1. Mutalik, V. K. et al. Precise and reliable gene expression via standard transcription and translation initiation elements. Nat. Methods 10, 354–360 (2013).

2. Bervoets, I. & Charlier, D. Diversity, versatility and complexity of bacterial gene regulation mechanisms: opportunities and drawbacks for applications in synthetic biology. FEMS Microbiol. Rev. 43, 304–339 (2019).

3. Shi, Y. Mechanistic insights into precursor messenger RNA splicing by the spliceosome. Nat. Rev. Mol. Cell Biol. 18, 655–670 (2017).

4. Lalanne, J.-B. et al. Evolutionary Convergence of Pathway-Specific Enzyme Expression Stoichiometry. Cell 173, 749–761.e38 (2018).

5. Dar, D. et al. Term-seq reveals abundant ribo-regulation of antibiotics resistance in bacteria. Science 352, aad9822 (2016).

6. Chen, Y.-J. et al. Characterization of 582 natural and synthetic terminators and quantification of their design constraints. Nat. Methods 10, 659–664 (2013).

7. Nielsen, A. A. K. et al. Genetic circuit design automation. Science 352, aac7341 (2016).

8. Park, Y., Espah Borujeni, A., Gorochowski, T. E., Shin, J. & Voigt, C. A. Precision design of stable genetic circuits carried in highly-insulated E. coli genomic landing pads. Mol. Syst. Biol. 16, e9584 (2020).

9. Green, A. A. et al. Complex cellular logic computation using ribocomputing devices. Nature 548, 117–121 (2017).

10. McCarty, N. S., Graham, A. E., Studená, L. & Ledesma-Amaro, R. Multiplexed CRISPR technologies for gene editing and transcriptional regulation. Nat. Commun. 11, 1281 (2020).

11. Cambray, G. et al. Measurement and modeling of intrinsic transcription terminators. Nucleic Acids Res. 44, 7006 (2016).

12. Li, R., Zhang, Q., Li, J. & Shi, H. Effects of cooperation between translating ribosome and RNA polymerase on termination efficiency of the Rho-independent terminator. Nucleic Acids Res. 44, 2554–2563 (2016).

13. Gorochowski, T. E. et al. Genetic circuit characterization and debugging using RNA-seq. Mol. Syst. Biol. 13, 952 (2017).

14. Hudson, A. J. & Wieden, H.-J. Rapid generation of sequence-diverse terminator libraries and their parameterization using quantitative Term-Seq. Synthetic Biology vol. 4 (2019).

15. He, Z. et al. Evaluating Terminator Strength Based on Differentiating Effects on Transcription and Translation. Chembiochem 21, 2067–2072 (2020).

16. Du, L., Gao, R. & Forster, A. C. Engineering multigene expression in vitro and in vivo with small terminators for T7 RNA polymerase. Biotechnol. Bioeng. 104, 1189–1196 (2009).

17. Mairhofer, J., Wittwer, A., Cserjan-Puschmann, M. & Striedner, G. Preventing T7 RNA polymerase read-through transcription-A synthetic termination signal capable of improving bioprocess stability. ACS Synth. Biol. 4, 265–273 (2015).

18. Schwarz-Schilling, M. et al. Correction to Optimized Assembly of a Multifunctional RNA-Protein Nanostructure in a Cell-Free Gene Expression System. Nano Lett. 19, 4812 (2019).

19. Liang, X., Li, C., Wang, W. & Li, Q. Integrating T7 RNA Polymerase and Its Cognate Transcriptional Units for a Host-Independent and Stable Expression System in Single Plasmid. ACS Synth. Biol. 7, 1424–1435 (2018).

20. Schaffter, S. W. & Schulman, R. Building in vitro transcriptional regulatory networks by successively integrating multiple functional circuit modules. Nat. Chem. 11, 829–838 (2019).

21. Liu, C. C., Jewett, M. C., Chin, J. W. & Voigt, C. A. Toward an orthogonal central dogma. Nat. Chem. Biol. 14, 103–106 (2018).

22. Wang, W. et al. Bacteriophage T7 transcription system: an enabling tool in synthetic biology. Biotechnol. Adv. 36, 2129–2137 (2018).

23. Jack, B. R., Boutz, D. R., Paff, M. L., Smith, B. L. & Wilke, C. O. Transcript degradation and codon usage regulate gene expression in a lytic phage†. Virus Evolution vol. 5 (2019).

24. Lyakhov, D. L. et al. Pausing and termination by bacteriophage T7 RNA polymerase. J. Mol. Biol. 280, 201–213 (1998).

25. Macdonald, L. E., Durbin, R. K., Dunn, J. J. & McAllister, W. T. Characterization of two types of termination signal for bacteriophage T7 RNA polymerase. J. Mol. Biol. 238, 145–158 (1994).

26. Porrua, O., Boudvillain, M. & Libri, D. Transcription Termination: Variations on Common Themes. Trends Genet. 32, 508–522 (2016).

27. Ju, X., Li, D. & Liu, S. Full-length RNA profiling reveals pervasive bidirectional transcription terminators in bacteria. Nat Microbiol 4, 1907–1918 (2019).

28. Chen, L. J. & Orozco, E. M., Jr. Recognition of prokaryotic transcription terminators by spinach chloroplast RNA polymerase. Nucleic Acids Res. 16, 8411–8431 (1988).

29. Casini, A. et al. R2oDNA designer: computational design of biologically neutral synthetic DNA sequences. ACS Synth. Biol. 3, 525–528 (2014).

30. Gruber, A. R., Lorenz, R., Bernhart, S. H., Neuböck, R. & Hofacker, I. L. The Vienna RNA websuite. Nucleic Acids Res. 36, W70–4 (2008).

31. Carr, S. B., Beal, J. & Densmore, D. M. Reducing DNA context dependence in bacterial promoters. PLoS One 12, e0176013 (2017).

32. Liu, S. S., Hockenberry, A. J., Lancichinetti, A., Jewett, M. C. & Amaral, L. A. N. NullSeq: A Tool for Generating Random Coding Sequences with Desired Amino Acid and GC Contents. PLOS Computational Biology vol. 12 e1005184 (2016).

33. Woodruff, L. B. A. et al. Registry in a tube: multiplexed pools of retrievable parts for genetic design space exploration. Nucleic Acids Res. 45, 1567–1568 (2017).

34. Potapov, V. et al. Comprehensive Profiling of Four Base Overhang Ligation Fidelity by T4 DNA Ligase and Application to DNA Assembly. ACS Synth. Biol. 7, 2665–2674 (2018).

35. Patrick, W. M., Firth, A. E. & Blackburn, J. M. User-friendly algorithms for estimating completeness and diversity in randomized protein-encoding libraries. Protein Eng. 16, 451–457 (2003).

36. Gorochowski, T. E. et al. Absolute quantification of translational regulation and burden using combined sequencing approaches. Mol. Syst. Biol. 15, e8719 (2019).

37. Garalde, D. R. et al. Highly parallel direct RNA sequencing on an array of nanopores. Nature Methods vol. 15 201–206 (2018).

38. Depledge, D. P. et al. Direct RNA sequencing on nanopore arrays redefines the transcriptional complexity of a viral pathogen. Nat. Commun. 10, 754 (2019).

39. Li, H. Minimap2: pairwise alignment for nucleotide sequences. Bioinformatics 34, 3094–3100 (2018).

40. Altschul, S. F., Gish, W., Miller, W., Myers, E. W. & Lipman, D. J. Basic local alignment search tool. J. Mol. Biol. 215, 403–410 (1990).

41. Li, H. et al. The Sequence Alignment/Map format and SAMtools. Bioinformatics 25, 2078–2079 (2009).

42. Stryer, L., Berg, J., Tymoczko, J. & Gatto, G. Biochemistry. (WH Freeman, 2019).

43. Loose, M., Malla, S. & Stout, M. Real-time selective sequencing using nanopore technology. Nat. Methods 13, 751–754 (2016).

44. Gilliot, P.-A. & Gorochowski, T. E. Sequencing enabling design and learning in synthetic biology. Curr. Opin. Chem. Biol. 58, 54–62 (2020).

45. Salis, H. M., Mirsky, E. A. & Voigt, C. A. Automated design of synthetic ribosome binding sites to control protein expression. Nat. Biotechnol. 27, 946–950 (2009).

46. Valeri, J. A. et al. Sequence-to-function deep learning frameworks for engineered riboregulators. Nat. Commun. 11, 5058 (2020).

47. Kotopka, B. J. & Smolke, C. D. Model-driven generation of artificial yeast promoters. Nat. Commun. 11, 2113 (2020).

48. Espah Borujeni, A., Zhang, J., Doosthosseini, H., Nielsen, A. A. K. & Voigt, C. A. Genetic circuit characterization by inferring RNA polymerase movement and ribosome usage. Nat. Commun. 11, 5001 (2020).

49. Bartoli, V., di Bernardo, M. & Gorochowski, T. E. Self-adaptive biosystems through tunable genetic parts and circuits. Current Opinion in Systems Biology 24, 78–85 (2020).

50. Vaser, R., Sović, I., Nagarajan, N. & Šikić, M. Fast and accurate de novo genome assembly from long uncorrected reads. Genome Res. 27, 737–746 (2017).

51. Tatusova, T. A. & Madden, T. L. BLAST 2 Sequences, a new tool for comparing protein and nucleotide sequences. FEMS Microbiol. Lett. 174, 247–250 (1999).

52. Baig, H. et al. Synthetic biology open language visual (SBOL visual) version 2.2. J. Integr. Bioinform. (2020) doi:10.1515/jib-2020-0014.

53. Bartoli, V., Dixon, D. O. R. & Gorochowski, T. E. Automated Visualization of Genetic Designs Using DNAplotlib. Methods Mol. Biol. 1772, 399–409 (2018).

