## Supplementary Information for "Massively parallel characterization of engineered transcript isoforms using direct RNA sequencing"

| **Supplementary Notes** | | **Page** |
|---|---|---|
| Supplementary Note 1: | Library coverage calculation | 2 |
| Supplementary Note 2: | Modelling direct RNA sequencing | 3 |
| **Supplementary Figures** | | |
| Supplementary Figure 1: | Analysis of assembled and sequenced library | 5 |
| Supplementary Figure 2: | Design of library used to optimize demultiplexing | 6 |
| Supplementary Figure 3: | Fitting model to direct RNA sequencing data | 7 |
| Supplementary Figure 4: | Deviation between observed and actual termination efficiencies | 8 |
| Supplementary Figure 5: | Polyadenylation efficiencies | 9 |
| Supplementary Figure 6: | Comparison of termination efficiencies calculated from biological replicates | 10 |
| Supplementary Figure 7: | Analysis of possible predictors of termination efficiency | 11 |
| **Supplementary Tables** | | |
| Supplementary Table 1: | Oligonucleotide sequences | 12 |
| **Supplementary References** | | 15 |

**Supplementary Note 1: Library coverage calculation.**

We estimated library coverage using the approach presented by Patrick *et al.*[1] to calculate the expected number of distinct sequences in a library chosen at random from a set of sequence variants. Given a pooled library containing $L$ sequences, and a set of $V$ equiprobable variants, let $v_i$ be one of the possible variants. Since the variants are equiprobable, the mean number of occurrences of $v_i$ in $L$ is

$$\lambda = L \,/\, V. \tag{S1}$$

For $\lambda \ll L$ (i.e. $V \gg 1$), the actual number of occurrences of $v_i$ in $L$ is essentially independent of the number of occurrences of any other variant $v_j$ where $j \neq i$, and therefore well-approximated by a Poisson distribution

$$P(x) = \frac{e^{-\lambda}\lambda^{x}}{x!}, \tag{S2}$$

where $P(x)$ gives the probability that $v_i$ occurs exactly $x$ times in the library. The probability that $v_i$ occurs at least once is given by $1 - P(0) = 1 - e^{-\lambda} = 1 - e^{-L/V}$. Therefore, the number of distinct variants expected in the library is given by

$$C \approx V(1 - e^{-L/V}), \tag{S3}$$

and the fractional completeness of the library is

$$F = \frac{C}{V} \approx 1 - e^{-L/V}. \tag{S4}$$

The library size required for fractional completeness $F$ is therefore

$$L \approx -V\ln(1 - F). \tag{S5}$$

In our case, $V$ = 1183 variants and we require a fractional completeness of $F > 1 - \frac{1}{1183} = 0.99915$ to ensure with high probability the representation of all variants in the library. This necessitates a library size of at least $L \approx -V\ln(1 - 0.99915) = 8364$. To achieve this, we performed a transformation protocol that used 10 large trays with approximately 50,000 transformants per tray (**Methods**), resulting in $L \approx 500000$.

### Supplementary Note 2: Modelling direct RNA sequencing

We developed a simple probabilistic model to capture the key processes impacting the reads recovered from a direct RNA sequencing (dRNA-seq) run. The following figure provides an overview of the major steps.

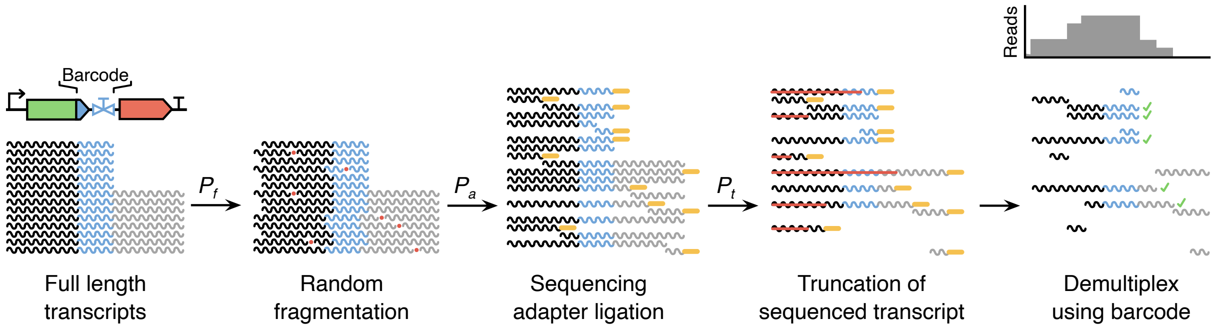


**Overview of the direct RNA sequencing model.** Reads are denoted by squiggles that are color coded to show core regions (e.g. blue region is the intrinsic barcode). Red dots show points of random fragmentation, orange oblongs represent sequencing adapters attached to only the 3' end of an RNA molecule, and green ticks denote reads that contain a complete barcode sequenced and which are used to generate a read depth profile. $P_f$, $P_a$, and $P_t$ are probabilities that reads are selected for each of the modification steps (i.e. random fragmentation, adapter ligation, and truncation, respectively).

We begin by assuming that all starting RNA transcripts are all full length corresponding to either an isoform that terminates at the transcriptional valve or reads through to the end of the construct. Then, reads are chosen with probability $P_f$ to become fragmented once at a random location along their length. This step captures the inevitable fragmentation that occurs when extracting and purifying an RNA sample. Next, a sequencing adapter is attached to each transcript or part of a fragmented RNA with probability $P_a$ and only molecules with an adapter attached are taken forward for sequencing. Sequenced molecules are then chosen with probability $P_t$ for possible truncation at a random position along the sequence. This step captures possible further fragmentation of the RNA during sequencing library preparation whereby only the fragment containing the adapter is sequenced, or possible truncation of reads due to premature termination during the sequencing of a molecule. Finally, we take the sequenced reads and filter out any that do not contain a complete transcriptional valve design (i.e. intrinsic barcode). Reads without a full barcode cannot be uniquely identified and so the reads are removed during the demultiplexing step. Reads that make it through these steps are then be used to generate a read depth profile.

To demonstrate the model's ability to capture real read depth profiles, we made use of the RNA Control Strand (CS) that is externally 'spiked-in' to all dRNA-seq runs for Quality Control

(QC) purposes. Crucially, the RNA CS is a single known sequence unlike any other in our library and only consists of full-length RNA molecules. Fitting our model to dRNA-seq data from the two biological replicates, we found that parameter values of $P_f$ = 0.1, $P_a$ = 0.77 and $P_t$ = 0.7 enabled a close fit in both cases, with only minor deviations at 5' and 3' ends of the RNA CS sequence (**Supplementary Figure 3A**). We also assumed the presence of an intrinsic barcode in the center of the RNA CS sequence and found that our model could also accurately predict read depth profiles recovered after demultiplexing of the real dRNA-seq data (**Supplementary Figure 3B**). This suggests that the read distribution that is generated by the model fits closely fits that recovered from sequencing and allowed us to further explore how well the observed read depth profiles matched the ground truth.

To explore this further, we used the model with parameters fitting to the real dRNA-seq data to simulate the sequencing process on synthetically generated transcripts for a hypothetical set of transcriptional valves with termination efficiencies varying between 0 and 1. By comparing the actual termination efficiency of each valve with the observed termination efficiency measured from the generated read depth profiles, we found a slight over estimation in $T_e$ (**Supplementary Figure 4**). To ensure this didn't bias our measurements for the data from the real transcriptional valves, this deviation was corrected for by subtracting the calculated error from the observed termination efficiency seen the model simulations to give a final $T_e$ value.

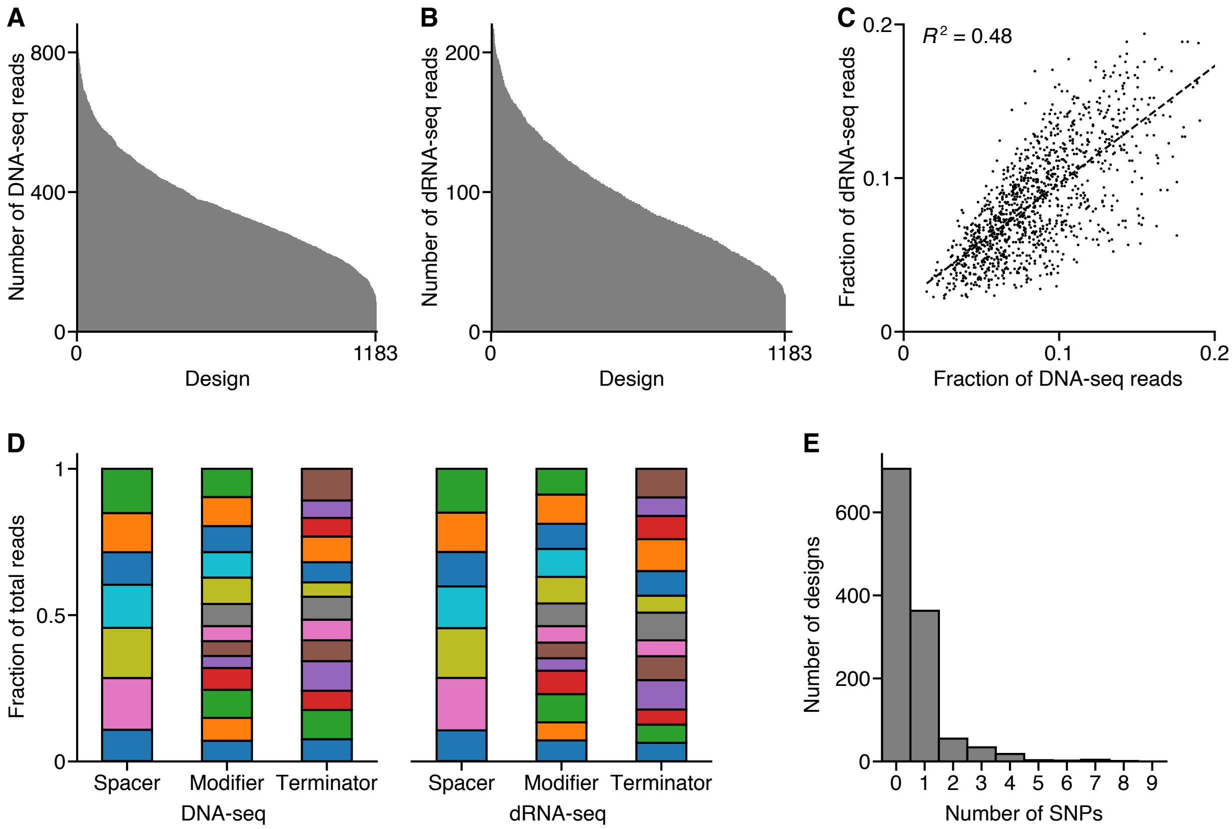


**Supplementary Figure 1: Analysis of assembled and sequenced library.** (**A**) Number of DNA-seq reads for each design, ordered by number of reads. (**B**) Number of dRNA-seq reads for each design, ordered by number of reads. (**C**) Comparison of frequency of DNA-seq and dRNA-seq reads. Each point corresponds to a single design and $R^2$ is the square of the Pearson correlation coefficient. (**D**) Frequency of each part in the DNA-seq (left) and dRNA-seq (right) data. Part and design frequencies were calculated relative to the total number of annotated sequencing reads. (**E**) Number of single nucleotide polymorphisms (SNP) per design.

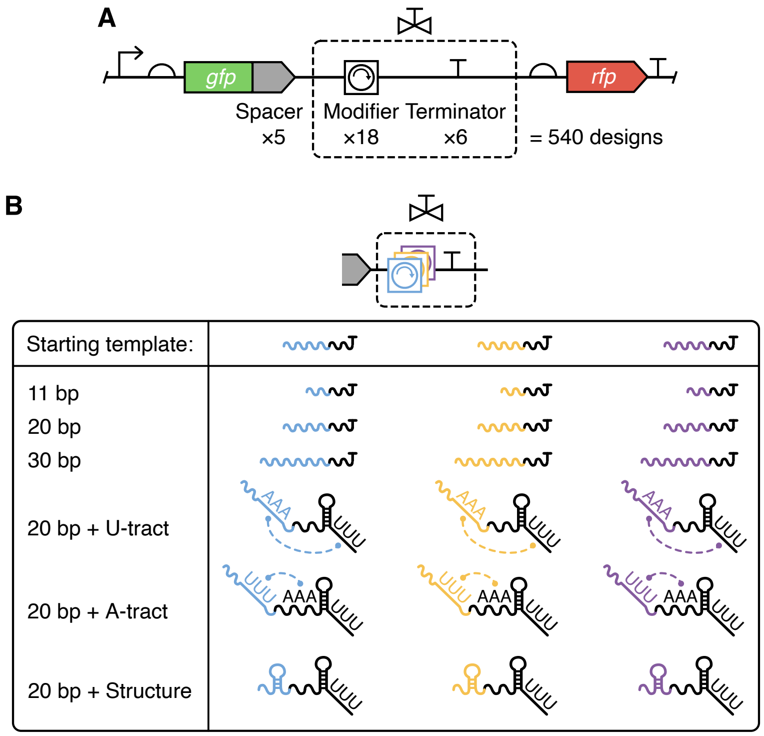


**Supplementary Figure 2: Design of library used to optimize demultiplexing.** (**A**) The library consists of 5 spacers (S1–S5), 18 modifiers (all parts with references beginning with M1–M3) and 6 terminators (T2–T7), resulting in 540 unique designs. For part sequences see **Supplementary Table 1**. (**B**) Modifiers were based upon 3 random starting template sequences, represented by different colored subsequences. From each template sequence 6 variants were made, each containing different proportions of the template sequence indicated by the number of base pairs: 11 bp sub-sequence, 20 bp sub-sequence, full 30 bp sequence, a 20 bp sub-sequence with U-tract interactor motif, a 20 bp sub-sequence with A-tract interactor motif, a 20 bp sub-sequence with structural motif.

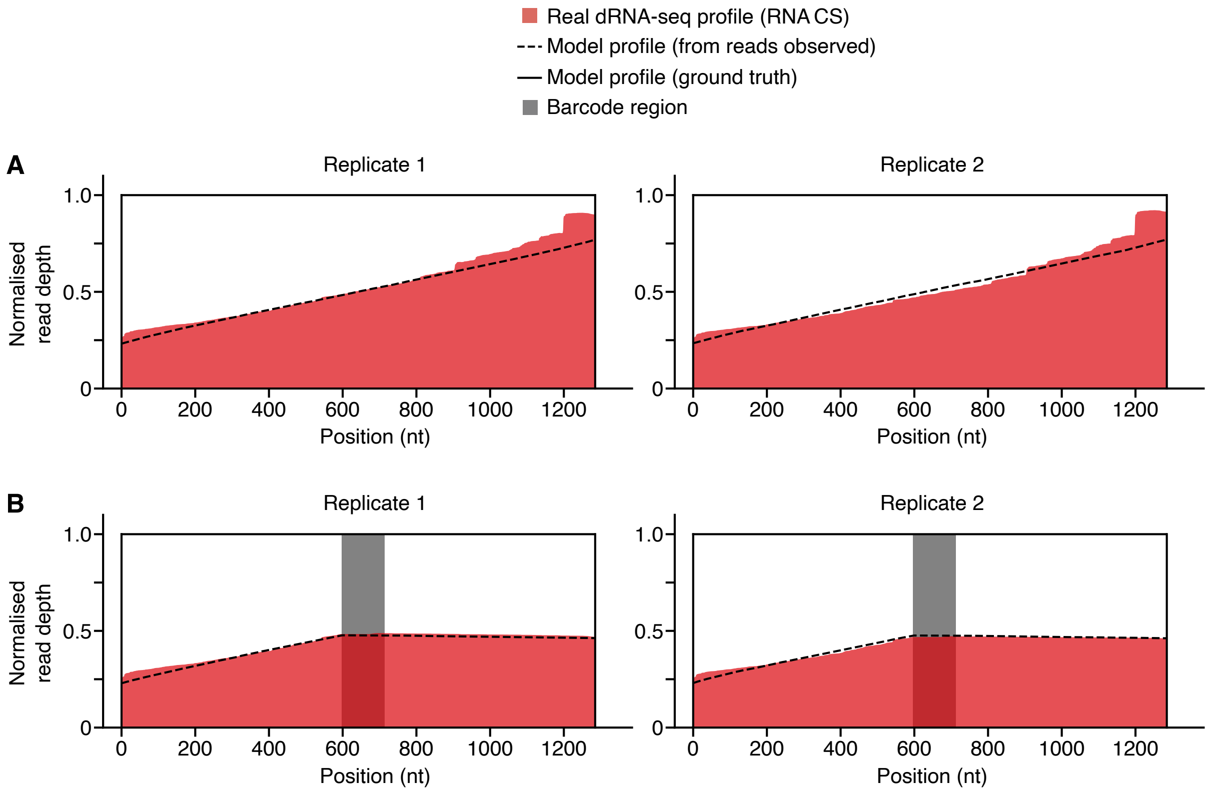


**Supplementary Figure 3: Fitting model to direct RNA sequencing data.** (**A**) Read depth profiles shown for all reads mapping to the RNA CS sequence for two dRNA-seq biological replicates (filled red) and fitted dRNA-seq model used to simulate the processing of 100,000 synthetic reads where $P_f$ = 0.1, $P_a$ = 0.77, $P_t$ = 0.7 (dashed black line for observed profile, solid black line for the model ground truth). (**B**) Read depth profiles for reads that map to the grey ‘intrinsic barcode’ for the real dRNA-seq data and fitted model.

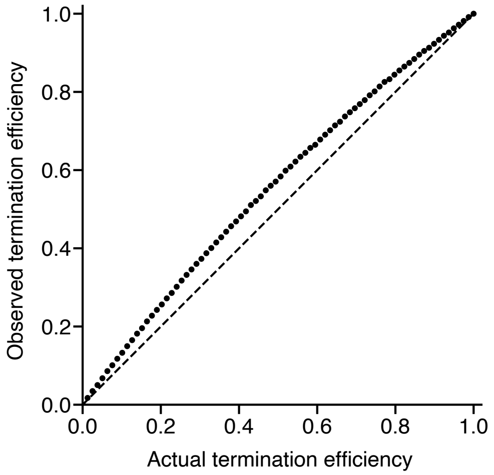


**Supplementary Figure 4: Deviation between observed and actual termination efficiencies.** Each point denotes a model simulation based on 100,000 artificially generated reads for transcriptional valves with varying termination efficiencies (**Supplementary Note 2**). Dashed line shows $y = x$.

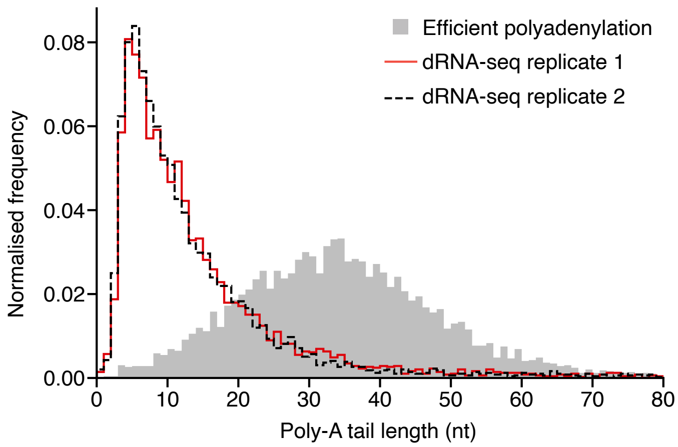


**Supplementary Figure 5: Polyadenylation efficiencies.** Histograms showing the varying lengths of RNA poly-A tail lengths for the two biological replicates analyzed in this work (dashed black and solid red lines) and another dRNA-seq sample where efficient polyadenylation was observed (grey filled histogram).

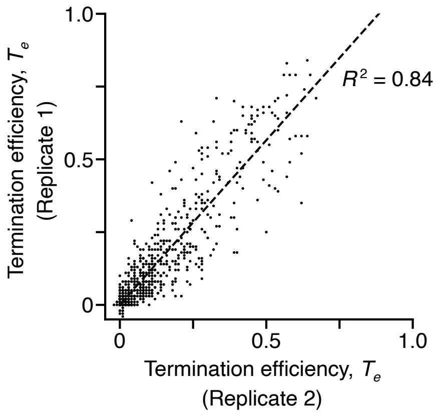


**Supplementary Figure 6: Comparison of termination efficiencies calculated from biological replicates.** Each point represents a single transcriptional valve design and dotted line shows the linear regression. $R^2$ is the square of the Pearson correlation coefficient.

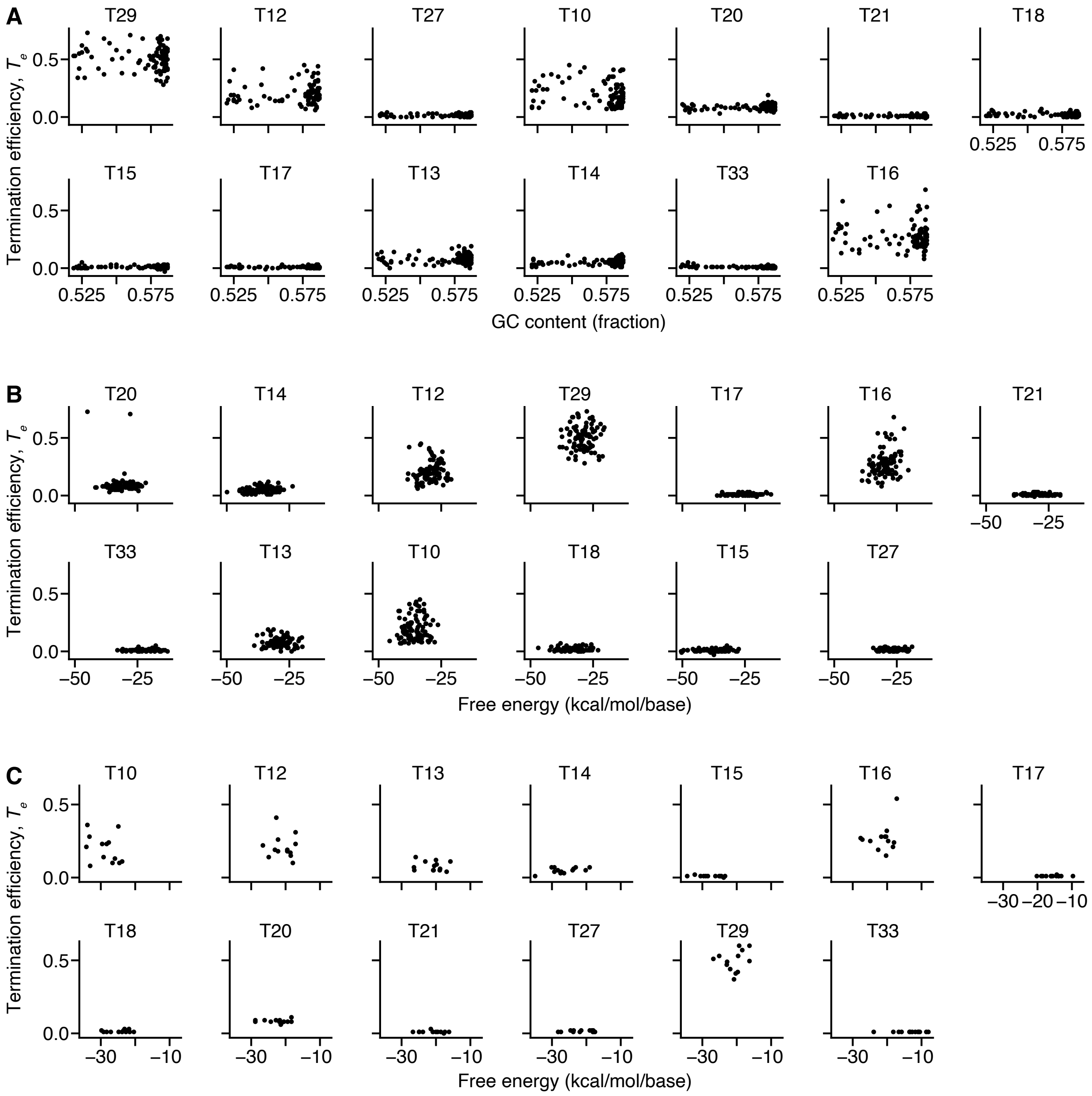


**Supplementary Figure 7: Analysis of possible predictors of termination efficiency.** (**A**) Scatter plot for each terminator showing $T_e$ against percentage GC content of each design. Calculation based on 80 nucleotides upstream of 3'-end of design. (**B**) Scatter plot for each terminator showing $T_e$ against the thermodynamic minimum free energy of each design. Calculation based on 120 nt upstream of 3'-end of design. (**C**) Scatter plot for each valve showing $T_e$ against the thermodynamic minimum free energy of each valve sequence.

### Supplementary Table 1: Oligonucleotide sequences

| ID | Forward strand oligonucleotide sequence | Reverse strand oligonucleotide sequence |
|---|---|---|
| pT7 | CTAATACGACTCACTATAGGGAGAG | CTAGCTCTCCCTATAGTGAGTCGTATTAGACGT |
| S10 | AATTCCTGTGTACCGGGAACCAGCCAGACTACACAGGGTAA | GCTCTTACCCTGTGTAGTCTGGCTGGTTCCCGGTACACAGG |
| S16 | AATTCGTGCAGAGACAAGCGTTTGGGGCACCAGCACAGTAA | GCTCTTACTGTGCTGGTGCCCCAAACGCTTGTCTCTGCACG |
| S18 | AATTCTTCAAAGCTACGAGCGCTAGAGATGTGAGACCCTAA | GCTCTTAGGGTCTCACATCTCTAGCGCTCGTAGCTTTGAAG |
| S19 | AATTCCTAATTATGTCTCAAAAGCTCGAAGATTACACCTAA | GCTCTTAGGTGTAATCTTCGAGCTTTTGAGACATAATTAGG |
| S20 | AATTCTTGTCGCTAAAGAAACCTTTCCCAATTAATACATAA | GCTCTTATGTATTAATTGGGAAAGGTTTCTTTAGCGACAAG |
| S21 | AATTCGGAATCGCTGATCTACAGAACGGTCCTTATGGGTAA | GCTCTTACCCATAAGGACCGTTCTGTAGATCAGCGATTCCG |
| S22 | AATTCATCACTCACACATCGCTCGAGATCGGTACGGGGTAA | GCTCTTACCCCGTACCGATCTCGAGCGATGTGTGAGTGATG |
| M10 | GAGCTTTCTCCGAAGTGTAGTAAAAAAATAAAAA | GGCATTTTTATTTTTTTACTACACTTCGGAGAAA |
| M11 | GAGCGATTACAGAAGCGTGGTATTTTTTATTTTT | GGCAAAAATAAAAAATACCACGCTTCTGTAATC |
| M12 | GAGCCAGGAACTTATCAATAGTCGCCCGAAAGGG | GGCACCCTTTCGGGCGACTATTGATAAGTTCCTG |
| M13 | GAGCCCTATTTACCTCAGT | GGCAACTGAGGTAAATAGG |
| M14 | GAGCTAGACAGTAATACCC | GGCAGGGTATTACTGTCTA |
| M15 | GAGCCTATCTGGTGCTACA | GGCATGTAGCACCAGATAG |
| M16 | GAGCTTATCGGTTACCAGA | GGCATCTGGTAACCGATAA |
| M17 | GAGCGTATCCAGACTTATTGAGGTTTACGCACTA | GGCATAGTGCGTAAACCTCAATAAGTCTGGATAC |
| M18 | GAGCATTCGCTGAGAGTTACACGATACTGACTAT | GGCAATAGTCAGTATCGTGTAACTCTCAGCGAAT |
| M19 | GAGCTTGAAATCGGATACTTCCTGAACTGCGAAT | GGCAATTCGCAGTTCAGGAAGTATCCGATTTCAA |
| M20 | GAGCATAGACTTTCGTGGATTATTACCTTACAACTGATAGGACGGACTC | GGCAGAGTCCGTCCTATCAGTTGTAAGGTAATAATCCACGAAAGTCTAT |
| M21 | GAGCATAGCCGAGATTATCCACCAGCAACAGTTCGTTATTGTAGTGATT | GGCAAATCACTACAATAACGAACTGTTGCTGGTGGATAATCTCGGCTAT |
| M22 | GAGCAAGGCGTGACTACAACCAATCTTCTATTCTGCGAGAGTAAAGTTT | GGCAAAACTTTACTCTCGCAGAATAGAAGATTGGTTGTAGTCACGCCTT |
| T10 | TGCCGCTGATGCCAGAAAGGGTCCTGAATTTCAGGGCCCTTTTTTTACATGGATTGA | CTAGTCAATCCATGTAAAAAAAGGGCCCTGAAATTCAGGACCCTTTCTGGCATCAGC |

| | | |
|---|---|---|
| T12 | TGCCACTGATTTTTAAGGCGACTGATGAGTCGCCTTTTTTTTGTCTA | CTAGTAGACAAAAAAAAGGCGACTCATCAGTCGCCTTAAAAATCAGT |
| T13 | TGCCAGTTAACCAAAAAGGGGGGATTTTATCTCCCCTTTAATTTTTCCTA | CTAGTAGGAAAAATTAAAGGGGAGATAAAATCCCCCCTTTTTGGTTAACT |
| T14 | TGCCCGTGTTCCTGAACGCCCGCATATGCGGGCGTTTTGCTTTTTGA | CTAGTCAAAAAGCAAAACGCCCGCATATGCGGGCGTTCAGGAACACG |
| T15 | TGCCTCTGAATGCGTGCCCATTCCTGACGGAATGGGCATTTCTGCGCAA | CTAGTTGCGCAGAAATGCCCATTCCGTCAGGAATGGGCACGCATTCAGA |
| T16 | TGCCGTTATTAAATAGCCTGCCATCTGGCAGGCTTTTTTTATCGA | CTAGTCGATAAAAAAAGCCTGCCAGATGGCAGGCTATTTAATAAC |
| T17 | TGCCCGTCTGCGTATGGAACGTGGTAACGGTTCTACTGAAGATTTA | CTAGTAAATCTTCAGTAGAACCGTTACCACGTTCCATACGCAGACG |
| T18 | TGCCTACTTCTTACTCGCCCATCTGCAACGGATGGGCGAATTTATACCCA | CTAGTGGGTATAAATTCGCCCATCCGTTGCAGATGGGCGAGTAAGAAGTA |
| T20 | TGCCCTGAAATATCCAGCGGATCAAGAAAATTCGTTGGATATTTTTTA | CTAGTAAAAAATATCCAACGAATTTTCTTGATCCGCTGGATATTTCAG |
| T21 | TGCCAAACACGTAGGCCTGATAAGCGAAGCGCATCAGGCAGTTTTGCGTA | CTAGTACGCAAAACTGCCTGATGCGCTTCGCTTATCAGGCCTACGTGTTT |
| T27 | TGCCTTTCAGCAAAAAACCCCTCAAGACCCGTTTAGAGGCCCCAAGGGGTTATGCTAGGA | CTAGTCCTAGCATAACCCCTTGGGGCCTCTAAACGGGTCTTGAGGGGTTTTTTGCTGAAA |
| T29 | TGCCCAGAAATCATCCTTAGCGAAAGCTAAGGATTTTTTTTATCTGAAA | CTAGTTTCAGATAAAAAAAATCCTTAGCTTTCGCTAAGGATGATTTCTG |
| T33 | TGCCCAGCGTTGAACCTACGACAGTCTCTTATTGACGAGTAAAGTGCTA | CTAGTAGCACTTTACTCGTCAATAAGAGACTGTCGTAGGTTCAACGCTG |
| S1 | AATTCGACTTTCACGTGAACCTGTTCCCAATATAA | GCTCTTATATTGGGAACAGGTTCACGTGAAAGTCG |
| S2 | AATTCAATGTGGAACTCTTCGCTCATGTAGAATAA | GCTCTTATTCTACATGAGCGAAGAGTTCCACATTG |
| S3 | AATTCGGTGCAGCGGAGAAAAGATTTGCTACCTAA | GCTCTTAGGTAGCAAATCTTTTCTCCGCTGCACCG |
| S4 | AATTCCTTGATATAAAACTTCCGGGAGTAGGATAA | GCTCTTATCCTACTCCCGGAAGTTTTATATCAAGG |
| S5 | AATTCCAAGAACTCGTTTTCCTATATGGCGTCTAA | GCTCTTAGACGCCATATAGGAAAACGAGTTCTTGG |
| M1N | GAGCTTTCTCCGAAGTGTAGTAAATAAAGCGTCC | GGCAGGACGCTTTATTTACTACACTTCGGAGAAA |
| M1A | GAGCTTTCTCCGAAGTGTAGTAAATTTTATTTTT | GGCAAAAATAAAATTTACTACACTTCGGAGAAA |
| M1U | GAGCTTTCTCCGAAGTGTAGTAAAAAAATAAAAA | GGCATTTTTATTTTTTTACTACACTTCGGAGAAA |
| M1S | GAGCTTTCTCCGAAGTGTAGTAAACCCGAAAGGG | GGCACCCTTTCGGGTTTACTACACTTCGGAGAAA |
| M1T | GAGCTTTCTCCGAAGTGTAGTAAA | GGCATTTACTACACTTCGGAGAAA |
| M1X | GAGCTTTCTCCGAAG | GGCACTTCGGAGAAA |
| M2N | GAGCAAGGACTTTCTCTACTGATTGTAAGACCGA | GGCATCGGTCTTACAATCAGTAGAGAAAGTCCTT |

| | | |
|---|---|---|
| M2A | GAGCAAGGACTTTCTCTACTGATTTTTTATTTTT | GGCAAAAAATAAAAAATCAGTAGAGAAAGTCCTT |
| M2U | GAGCAAGGACTTTCTCTACTGATTAAAATAAAAA | GGCATTTTTATTTTAATCAGTAGAGAAAGTCCTT |
| M2S | GAGCAAGGACTTTCTCTACTGATTCCCGAAAGGG | GGCACCCTTTCGGGAATCAGTAGAGAAAGTCCTT |
| M2T | GAGCAAGGACTTTCTCTACTGATT | GGCAAATCAGTAGAGAAAGTCCTT |
| M2X | GAGCAAGGACTTTCT | GGCAAGAAAGTCCTT |
| M3N | GAGCCAGGAACTTATCAATAGTCGTTGTGACACT | GGCAAGTGTCACAACGACTATTGATAAGTTCCTG |
| M3A | GAGCCAGGAACTTATCAATAGTCGTTTTATTTTT | GGCAAAAAATAAAACGACTATTGATAAGTTCCTG |
| M3U | GAGCCAGGAACTTATCAATAGTCGAAAATAAAAA | GGCATTTTTATTTTCGACTATTGATAAGTTCCTG |
| M3S | GAGCCAGGAACTTATCAATAGTCGCCCGAAAGGG | GGCACCCTTTCGGGCGACTATTGATAAGTTCCTG |
| M3T | GAGCCAGGAACTTATCAATAGTCG | GGCACGACTATTGATAAGTTCCTG |
| M3X | GAGCCAGGAACTTAT | GGCAATAAGTTCCTG |
| T2 | TGCCCGTAAAAACCCGCCGAAGCGGGTTTTTACGTAACA | CTAGTGTTACGTAAAAACCCGCTTCGGCGGGTTTTTACG |
| T3 | TGCCAGTAAAAACCCGCCGAAGCGGGTTTTTACGTAACA | CTAGTGTTACGTAAAAACCCGCTTCGGCGGGTTTTTACT |
| T4 | TGCCAAAAAAAACACCCTAACGGGTGTTTTTTTTTTTTA | CTAGTAAAAAAAAAAAACACCCGTTAGGGTGTTTTTTTT |
| T5 | TGCCAGAATTCAGTCAAAAGCCTCCGACCGGAGGCTTTTGACTATTACTACTAGA | CTAGTCTAGTAGTAATAGTCAAAAGCCTCCGGTCGGAGGCTTTTGACTGAATTCT |
| T6 | TGCCAGAATTCAGCCCGCCTAATGAGCGGGCTTTTTTTTACTAA | CTAGTTAGTAAAAAAAAGCCCGCTCATTAGGCGGGCTGAATTCT |
| T7 | TGCCAGAAAAGAGGCCTCCCGAAAGGGGGGCCTTTTTTCGTTTTA | CTAGTAAAACGAAAAAAGGCCCCCCTTTCGGGAGGCCTCTTTTCT |

### Supplementary References

[1] Patrick, Wayne M., Andrew E. Firth, and Jonathan M. Blackburn. User-Friendly Algorithms for Estimating Completeness and Diversity in Randomized Protein-Encoding Libraries. *Protein Engineering* **16** (6): 451–57 (2003).
